# Saxitoxin potentiates Zika virus-induced cell death in human neurons but not in neural progenitors and astrocytes

**DOI:** 10.1101/2023.05.22.541218

**Authors:** Leticia R. Q. Souza, Carolina da S. G. Pedrosa, Teresa Puig-Pijuan, Camila da Silva dos Santos, Gabriela Vitória, Luiza M. Higa, Amilcar Tanuri, Marília Zaluar P. Guimarães, Stevens Kastrup Rehen

## Abstract

The Zika virus (ZIKV) outbreak in Brazil between 2015 and 2016 was associated with an increased prevalence of severe congenital malformations, including microcephaly. Notably, the distribution of microcephaly cases was not uniform across the country, with a disproportionately higher incidence recorded in the Northeast region (NE). Our previous work demonstrated that saxitoxin (STX), a toxin ubiquitously present in the drinking water reservoirs of the NE, exacerbated the damaging effects of ZIKV on the developing brain. In the present study, we hypothesized that STX’s impact might vary among different neural cell types. Our experimental observations suggest that exposure to STX potentiates the neurotoxic effect of Zika Virus (ZIKV) on human neuronal cells. However, while ZIKV infection demonstrated severe impacts on astrocytes and neural stem cells (NSCs), the addition of STX did not exacerbate these effects. We observed that neurons subjected to STX exposure were more prone to apoptosis and displayed a higher number of ZIKV-infected cells. These findings suggest that STX exacerbates the harmful effects of ZIKV on neurons, thereby providing a plausible explanation for the heightened severity of ZIKV-induced congenital malformations observed in Brazil’s NE. This study underscores the importance of understanding the interactive effects of environmental toxins and infectious pathogens on neural development, with potential implications for public health policies and interventions.

## INTRODUCTION

The Zika virus (ZIKV) has been conclusively linked to a range of congenital malformations, notably microcephaly, particularly when contracted in the first trimester of pregnancy. These outcomes are collectively referred to as Congenital Zika Syndrome (CZS) (1,2). The syndrome exhibits a broad array of abnormalities, spanning from fetal demise to craniofacial disproportion, ventriculomegaly, and arthrogryposis, among others (3–5).

The ZIKV epidemic that primarily affected Northeastern Brazil (NE) led to a markedly increased incidence of congenital malformations. Intriguingly, this region reported a significantly higher prevalence of such abnormalities relative to other parts of the world, leading researchers to suspect the presence of a co-factor that might account for this regional discrepancy in severity (6,7).

The NE region is characterized by frequent droughts that foster the proliferation of cyanobacteria, including *Cylindrospermopsis raciborskii*, a known producer of saxitoxin (STX) (8). STX is a member of the Paralytic Shellfish Toxins (PSTs) group, known to impact a variety of cell types, including those in the central nervous system (9,10). This toxin has been implicated in the disruption of normal neuronal development, and it can interfere with voltage-dependent sodium, potassium, and calcium channels, thereby disturbing electrical activity (11). The persistence of STX in water resources, particularly in economically disadvantaged areas, is a substantial public health concern. Strikingly, regions with high rates of microcephaly corresponded closely with areas of low-volume water reservoirs containing cyanobacterial toxins (12,13). The exceptional severity of ZIKV-induced malformations in the NE region, coupled with the endemic presence of STX, led to the hypothesis of a potential interaction between these two factors (14).

Our previous research showed an increased rate of cell death in the brains of neonatal mice born to mothers co-exposed to ZIKV and STX, and in human brain organoids under similar conditions. These findings implicated STX as a potential environmental factor exacerbating the impact of ZIKV (14). However, the cell types affected have yet to be determined. In the present study, we utilized a range of human neural cells to investigate this unresolved question. We aimed to identify the specific neural cell types that are impacted by concurrent ZIKV infection and STX exposure. This study seeks to enhance our understanding of the interaction between these two factors and their role in CZS.

## METHODS

### HUMAN INDUCED PLURIPOTENT STEM CELLS

All cell types used in the present work were derived from human induced pluripotent stem cells (hiPSCs) obtained after reprogramming somatic cells from three different healthy individuals. Two of them were derived from fibroblasts isolated from skin biopsies and the third one (GM23279A) is commercially available at the NIGMS Human Genetic Cell Repository, obtained and certified by the Coriell Institute for Medical Research cell bank. All in-house hiPSCs were reprogrammed using a previously published protocol (15) and were used in subsequent works by our group (16–21). All cells grown in monolayers used in this work were maintained at 37°C, in an atmosphere containing 5% CO_2_ and cultivated in the absence of any antibiotics.

### NEURAL STEM CELLS (NSCs)

Neural Stem Cells (NSCs) were generated from hiPSCs cells following the published Life Technologies protocol for “Induction of Neural Stem Cells from Human Pluripotent Stem Cells” (Publication number: MAN0008031). Briefly, the hiPSCs were maintained for 7 days in a medium containing Neurobasal (Thermo Fisher Scientific, 12348017) and 1% neural induction supplement (NIS – Life Technologies, a16477-01) which was changed every day on an adherent plate coated with Geltrex LDEV-Free (Thermo, A1413302). After induction, NSCs were grown in adherent 60 mm plates coated with Geltrex LDEV-Free and in NEM medium (Neural Expansion Medium), composed of 49% Neurobasal (Thermo Fisher Scientific, 12348017), 49% Advanced-DMEM (Thermo Fisher, 12634-010) and 2% neural induction supplement (Life Technologies, a16477-01). The medium was changed every 2 days and when 100% confluence was reached the cells were enzymatically dissociated with Accutase (Millipore, SCR005). For experiments, cells were plated onto 96 well plates at a density of 10,000 cells per well or in 24 well plates with glass coverslips at a density of 40,000 cells per well, both plates previously coated with Geltrex.

### GLIAL DIFFERENTIATION

Astroglial induction was based on the protocol previously described by Yan et al. (22,23). The protocol started with NSC plated at 1.25 x 10^6^ in a T-25 flask, previously coated with 1x Geltrex (Thermo, A1413302). On the day after plating, NEM medium was replaced by AIM medium (Astrocyte Induction Medium) composed of DMEM/F12 (Life Technologies, 11330-032), 1% Fetal Bovine Serum (FBS -Thermo, 12657029) and 1x N2 (Invitrogen, 17502001). The AIM medium was changed every 2 days for 21 days. During these first 21 days, cell cultures were passaged with Accutase when 90% confluence was reached and split 1:4. On the 14th day the coating of the flasks was made with a reduced concentration (0.5x) of Geltrex. After day 21, an astrocyte expansion medium consisting of DMEM/F12 with 10% FBS was added, and the cells passed to uncoated flasks. Cells were maintained for at least 6 weeks in this medium for further maturation. Glial cells between 6 and 20 weeks of maturation were used for the assays. For experiments, the cells were plated in 96 well plates with 2.5 x 10^3^ cells per well or in 24 well plates, containing glass coverslips, with a density of 2.5 x 10^4^ cells per well.

### NEURONAL DIFFERENTIATION

#### Human cortical neurons

The neuronal differentiation protocol was adapted from two previously described protocols (22). To obtain a mixed neuronal culture, 3.0 x 10^6^ NSCs were first seeded in a 100 mm plate, previously coated with 10 μg/mL laminin (Invitrogen, 23017-015) in NEM medium. The next day, the medium was changed to neuronal differentiation medium, which consists of Neurobasal (Life Technologies, 21103049), 1x B27 Supplement (Life Technologies, 17504044), 1x Glutamax (Life Technologies, 35050061), 1x non-essential amino acids (Life Technologies, 11140050) and 200 μM ascorbic acid (AA) (Sigma Aldrich, A4034). This medium was maintained until the end of the experiment, being changed every two days. On the 7th and 14th days, the cells were passed with Accutase to a plate coated with 10 μg/mL laminin and kept until the neurons completed 50 days. For experiments, the cells were plated in 96 well plates with 3 x 10^4^ cells per well or in 24 well plates, containing glass coverslips, with a density of 2 x 10^5^ cells per well.

#### Human sensory neurons

The sensory neuron differentiation protocol was made with some modifications from the protocol published by Guimarães et al. (24). NSCs (5 x 10^6^) were seeded in NEM medium onto 100 mm plates coated with 100 μg/mL polyornithine (PLO) (Sigma Aldrich, P3655) and 20 μg/mL laminin. Next day, the medium was changed to 3N for neuronal differentiation (DMEM-F12, Neurobasal Medium, 1x Glutamax, 0.5x N2 Supplement, 0.5x B27 Supplement, 0.5x non-essential amino acids, and 1:1000 β-mercaptoethanol (Sigma Aldrich, M3148)) supplemented with 10 ng/mL BDNF (R&D systems, 248-BD-025), 200 μM Ascorbic Acid (AA) (Sigma Aldrich, A4034), 10 ng/mL GDNF (R&D Systems, 212-GD-010), 10 ng/mL NGF (R&D Systems, 256-GF-100),10 ng/mL NT-3 (R&D Systems, 267-N3-025) and 0.5 mM cAMP (Sigma-Aldrich, D0260-100MG). Medium changes were performed twice a week. On day 45 cells were passaged with Accutase and incubated with the same neuronal medium supplemented with 10 μM iRock (Millipore) and plated for experiments. The cells were seeded in 96-well plates with 3 x 10^4^ cells per well or in 24-well plates, containing glass coverslips, with a density of 2 x 10^5^ cells per well.

### BRAIN ORGANOIDS

The production of organoids followed a protocol previously described by Goto-Silva et al. (25), in which hiPSCs were cultivated in mTeSR1 medium (StemCell Technologies, 05850) on Matrigel (BD Biosciences, CLS354277). When the colonies reached 70-80% confluency, hiPSCs were dissociated with Accutase and seeded at 9,000 cells/well into 96-well round bottom low attachment plates (Corning, 7007) in hiPSCs medium. The next day, the medium was replaced with an hESC medium, and the embryoid bodies (EBs) were cultured for 6 days in the hESC medium previously described in Lancaster et al. (2014) (26). On day 6 the EBs were transferred to 24-well flat bottom ultra-low attachment culture plates (Corning, 3473) containing Neural Induction Medium DMEM/F12, 1% Pen-Strep (Thermo Fisher Scientific, 15140122), 1% N-2, 1% GlutaMAX, 1% MEM-NEAA and 1 μg/mL heparin (Thermo Fisher Scientific, H3149-100KU) for 4 days. The organoids were coated with Matrigel for 1 hour at 37°C and 5% CO_2_ and returned to 24-well flat bottom ultra-low attachment plates. The organoids were maintained for 4 days in static culture in neural differentiation medium, DMEM/F12, and Neurobasal medium (1:1), 1% Pen-Strep, 0.5% N-2, 1% B-27 without vitamin A (Thermo Fisher Scientific, 17504044), 1% GlutaMAX, 1% MEM-NEAA, 0.035% 2-mercaptoethanol and 1:4000 insulin (Sigma Aldrich, I9278) and after that time were cultured in suspension in 6 wells plates, under constant stirring at 90 rpm on an orbital shaker. For this final step, 4-6 organoids were allocated per well containing 3 mL of neural differentiation medium with added vitamin A (only changing the B27 supplement to Thermo Fisher Scientific, 17504001). The medium was changed twice a week until completing 45 days of agitation when they were used for experiments.

### REPLICATION AND VIRAL INFECTION

The Asian viral strain (AS) ZIKV (Recife/Brazil, ZIKV PE/243, number: KX197192.1) was provided by Dr. Marli Tenório Cordeiro from Fundação Oswaldo Cruz/Centro de Pesquisas Aggeu Magalhães, Brazil. The virus was propagated in C6/36 *Aedes albopictus* cell line at a multiplicity of infection (MOI) of 0.01 and cultured for 6 days in Leibovitz’s L-15 medium (Thermo Fisher Scientific, 11415064) supplemented with 0.3% tryptose phosphate broth (Sigma-Aldrich, T4532), 2 mM L-glutamine (Thermo Fisher Scientific, 25030081) and 1x MEM non-essential amino acids and 2% FBS. ZIKV titers were determined by conventional plaque assay.

NSCs, astrocytes, and neurons were treated with saxitoxin for 1 week at 6 ng/L (neurons only) or 12 ng/L (all cell types). After that, cells were washed once with 1x PBS and incubated with the viral inoculum for 2 hours at an MOI = 0.5. Infection controls were incubated with the supernatant of uninfected C6/36 *Aedes albopictus* cell line (MOCK). After this time, the viral inoculum was removed, and the cells were incubated for another 72 hours in their respective culture media containing saxitoxin. The organoids were infected for 2 hours at MOI 0.5 and then maintained for up to 13 days. Saxitoxin was added at a concentration of 12 ng/L immediately after the 2 hours of infection and added at each medium change throughout the 13 days.

### MTT CELL VIABILITY ASSAY

After 72 hours of infection, 10 μL of 0.5 mg/mL MTT (3-(4,5-dimetiltiazole-2yl)-2.5-diphenyl tetrazolium), diluted in DMEM/F12, was added per well and the cells were incubated for 3 hours at 37°C. Following this period, the MTT solution was removed and 100 μL of DMSO was added per well. Then, the absorbances at wavelengths of 560 nm and 690 nm were determined in the Tecan Infinite 200 PRO plate reader. The absorbance was corrected by calculating the difference between values obtained at 560 nm and the reference wavelength (690 nm). The relative viability was calculated considering the untreated uninfected group as 100% viability.

### IMMUNOFLUORESCENCE ANALYSIS

Cells were fixed in 4% PFA for 20 minutes at 37°C after 72 h. The cells were washed 3 times with 1x PBS and incubated for 2 hours with 5% goat serum (NGS -Sigma-Aldrich, G9023) in 1x PBS with 0.3% Triton (Sigma-Aldrich, T8787). Then, the primary antibodies [Non-structural protein 1 from ZIKV (NS1, 1:500, BioFront Technologies, BF1225-06;), Class III β-tubulin (TuJ3, 1:200, Sigma-Aldrich, t3952), MAP2 (1:200, Thermo Fischer Scientific PA517646), and TRPV1 (1:250, Abcam, ab3487) were diluted in the same solution and incubated overnight at 4°C. Next day, the coverslips were washed 3 times with PBS 1x, and incubated with secondary antibodies AlexaFluor 488 (1:400, Thermo Fischer Scientific A-11008 and A-11001) and AlexaFluor 647 (1:400, Thermo Fischer Scientific A31573 and A32728) diluted in PBS for 2 hours at room temperature after which they were counterstained with 300 nM 4’,6’-diamino-2-phenylindole (DAPI; 10236276001, Roche) for 5 minutes, the coverslips were rewashed with 1x PBS and milli-Q water and finally mounted with AquaMount (Polysciences, 18606-20). To assess cell death, the Click-iT™ TUNEL Alexa Fluor™ 594 Imaging Assay was used (Thermo Fisher Scientific, C10246) following the data sheet provided by the manufacturer. All images were acquired on a Leica TCS SP8 confocal microscope. Infection was quantified by counting the number of NS1 positive cells per field compared to the total number of cells (DAPI positive). Cell death was quantified by counting the number of tunnel-positive cells per field compared to the total number of cells (DAPI positive). At least 10 fields per coverslip (average of 2 coverslips per experimental group) were quantified and 3 independent experiments were performed. Quantification was obtained using the Cell Counter plugin of the ImageJ software.

### ISOTROPIC FRACTIONATION

ZIKV-infected organoids treated with saxitoxin were fixed and the nuclei of these organoids were obtained by isotropic fractionation following the protocol in Herculano and Lent (27). These nuclei were seeded in 384 plates coated with 0.1 mg/mL poly-L-lysine. Cell death was detected by the Click-iT™ TUNEL Alexa Fluor™ 594 Imaging Assay and staining was performed according to the manufacturer’s instructions. Antibodies NeuN (1:100, Millipore, MAB377) and TBR1 (1:200, Thermo Fisher Scientific, PA530971) were used to stain neuronal nuclei. The nuclei were labeled with 0.5 μg/mL of 40-6-diamino-2-phenylindole (DAPI) for 10 minutes. Nuclei were washed with 1x PBS, mounted with glycerol, and analyzed on an Operetta high-content imaging system with a 40x objective and high numerical apertures (NA) (PerkinElmer, USA). Data were analyzed using Harmony 5.1 high-content image analysis software (PerkinElmer, USA). Ten independent fields were evaluated from wells in triplicate per experimental condition using 3 organoids for each condition.

### STATISTICAL ANALYSIS

The data were expressed as the average ± standard error of the mean (SEM). Replicas values of the same conditions were submitted to the Grubbs test for outliers, in which values considered significantly discrepant (α=0.05) were disregarded. For the analyses of significance, One-Way ANOVA (Nonparametric or mixed) followed by Tukey’s multiple comparisons tests was used. The differences were considered statistically significant when p < 0.05.

## RESULTS

### Saxitoxin Augments Zika Virus-Induced Neuronal Loss and Cell Death in Brain Organoids

In our previous work, we demonstrated that the co-exposure of brain organoids to STX and ZIKV resulted in a 2.5-fold increase in cell death and enhanced ZIKV replication, while the proportion of neural progenitors was unaffected (16). However, the specific cellular subpopulations impacted by STX remained elusive. To address this, we adapted an isotropic fractionation method originally applied to the human brain (27) for quantifying cellular subpopulations within organoids. Through nuclear staining, we assessed the proportion of mature (NeuN+) and immature (TBR1+) neurons remaining after 13 days of exposure to STX and ZIKV in human brain organoids. We validated the method by confirming a 3.9-fold surge in cell death (TUNEL+ cells), aligning with the findings of Pedrosa et al. (14) (Figure 1A and 1B). Upon exposure to STX, ZIKV-infected immature (TBR1+) and mature (NeuN+) neurons were significantly impacted (Figure 1A, 1C, and 1D). Both categories of neurons exhibited a marked decline in their numbers when ZIKV-treated cells were compared with those co-exposed to ZIKV and STX. Specifically, the percentage of TBR1+ cells decreased by 5.6-fold and NeuN+ cells by 2.9-fold due to STX treatment. The remaining cellular populations within the ZIKV-infected organoids were not additionally affected by the addition of STX (data not shown).

**Figure 1.**
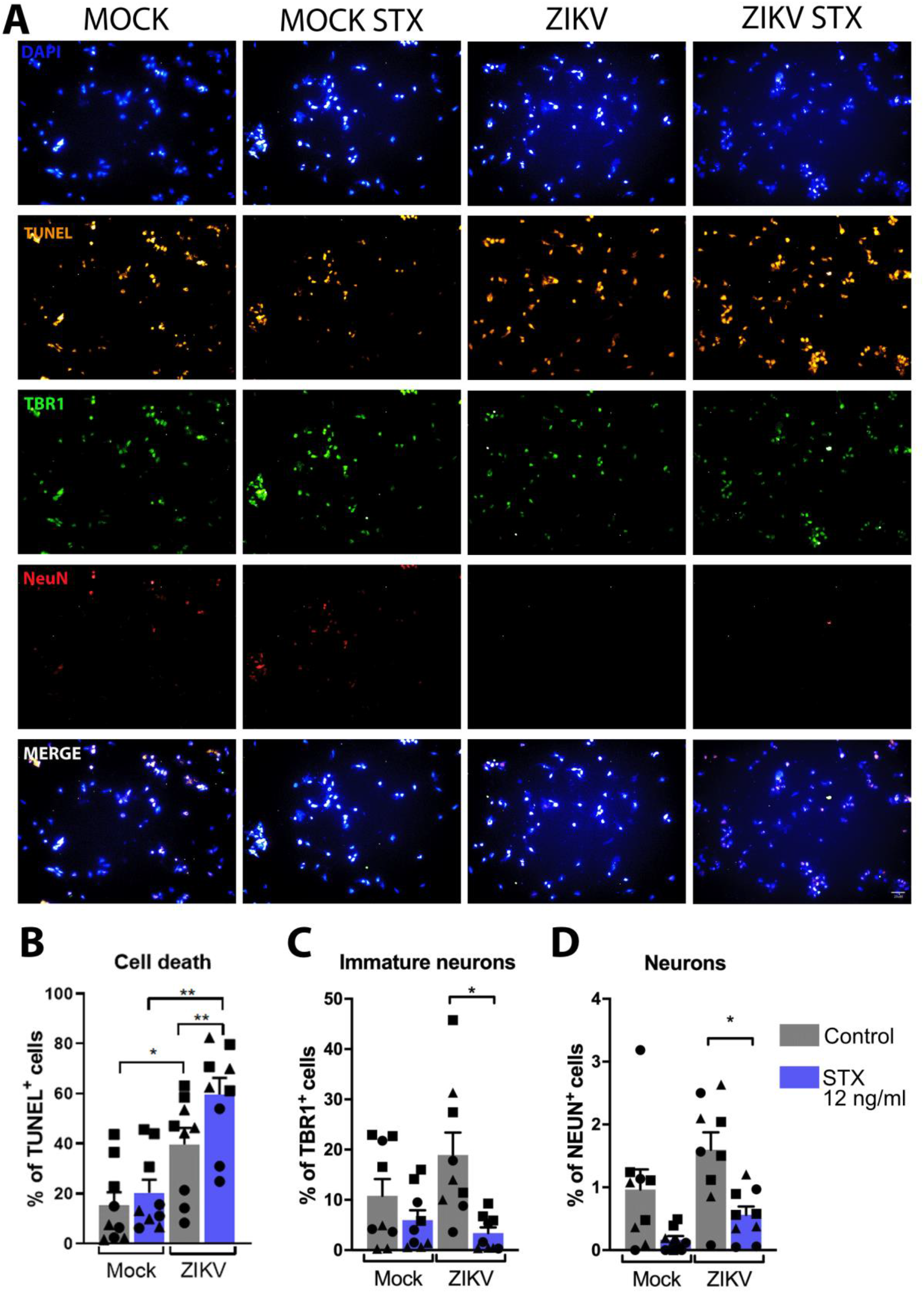
Neuronal populations within brain organoids exhibited a significant reduction following treatment with Saxitoxin and subsequent infection with the Zika virus. (A) Representative photomicrographs of nuclei of 55-days old brain organoid fractionated following STX treatment and ZIKV infection. The nuclei were labeled TUNEL (orange), TBR1 (green), and NeuN (red). Scale bar: 20 μm. Graphs of the percentage of labeled cells over total cells (DAPI+) in fractionated organoids after ZIKV infection and STX treatment. (B) TUNEL-positive cells. (C) TBR1-positive cells. (D) NeuN-positive cells. The experiments were performed with 3 different organoids, in 3 independent differentiations, 13 days after ZIKV infection and STX treatment. Data are presented as average ± SEM; *p<0.05 and **p<0.01. One-way ANOVA was followed by Tukey’s post-test.

### Saxitoxin exposure further exacerbates cell death in human cortical neurons induced by Zika virus infection

Subsequently, our goal was to discern if the concurrent exposure to saxitoxin and ZIKV would also impact individual neuronal populations or whether this effect was exclusive to more intricate 3D cellular models necessitating complex cell-cell interactions. To examine this, we differentiated human cortical neurons from neural stem cells (NSCs) over a 50-day period (Supplementary Figure 1A). These neurons were exposed to saxitoxin for seven days prior to ZIKV infection, followed by a further three-day re-incubation with saxitoxin. We observed in cortical neurons that cell death (TUNEL+) was not only induced by STX alone but was further amplified by ZIKV infection (Figures 2A and 2B). This was accompanied by a decline in cell viability as evidenced by the MTT assay (Figure 2C). Moreover, the proportion of ZIKV-infected cells (NS1+) had a 2.7-fold increase in the presence of STX (Figure 2D). These results are consistent with our earlier observations in 3D brain organoids, demonstrating similar effects in 2D cortical neurons.

**Figure 2.**
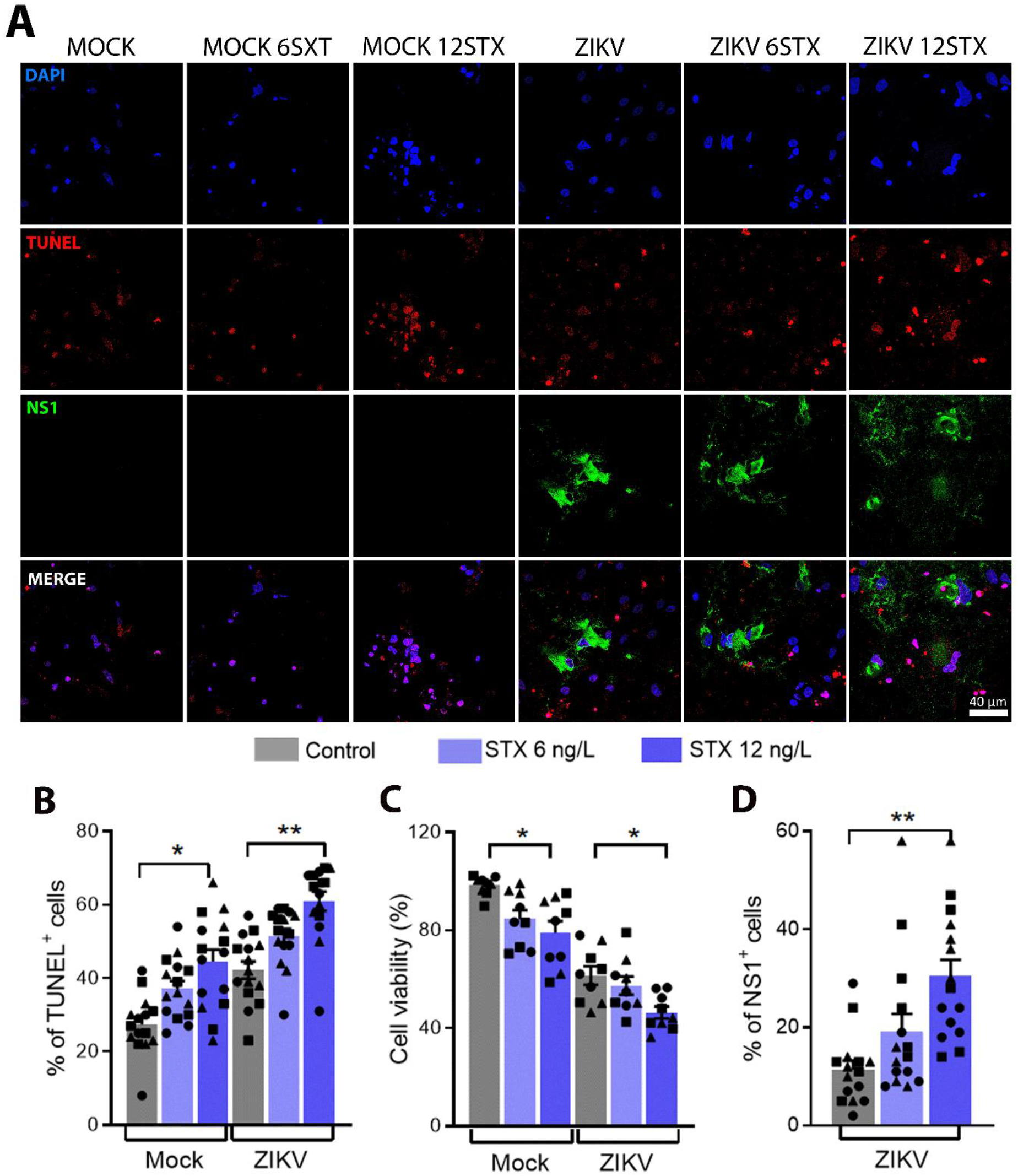
Saxitoxin increases cell death and the percentage of infection in ZIKV-infected hiPSC-derived cortical neurons. (A) Representative photomicrographs of TUNEL-positive cells (red) and NS1-positive cells (green) of untreated or STX-treated Mock and ZIKV-infected hiPSC-derived cortical neurons. Scale bar: 40 μm. (B) Percentage of TUNEL-positive cells per DAPI-positive cells. (C) MTT cell viability assay in hiPSC-derived cortical neurons following ZIKV infection and STX treatment. (D) Percentage of NS1-positive cells per DAPI-positive cells. Experiments were performed in triplicates, in 3 independent differentiations, using 2 cell lines per condition pre-treated with saxitoxin for 7 days, infected with ZIKV, and incubated with saxitoxin again for 3 days. Data are presented as average ± SEM; *p<0.05 and **p<0.01. One-way ANOVA followed by Tukey’s multiple comparisons test was performed.

### Effects of Zika Virus and Saxitoxin on Human Sensory Neurons

Congenital Zika Syndrome (SCZ) presents various sensory symptoms, including somatosensory ones (28,29). Despite numerous tests on hiPSC-derived neurons with ZIKV, there is yet no evidence that human sensory neurons can be infected. We previously reported a productive infection in hiPSC-derived peripheral neurons, but not specifically sensory ones (30).

To explore this, we exposed hiPSC-derived human sensory neurons, produced using our lab’s protocol (Supplementary Figure 1B) (24), to ZIKV. Upon confirming their susceptibility to infection, we investigated if STX could exacerbate ZIKV-induced damage in these neurons, similar to cortical neurons. Our findings revealed a threefold increase in cell death (TUNEL+) in sensory neurons subjected to 12 ng/L STX, compared to controls. When comparing ZIKV-infected neurons to those exposed to both ZIKV and STX, we observed a 2.1-fold increase in cell death (Figure 3A and 3B), coinciding with a decrease in viability (Figure 3C, MTT assay). Furthermore, we noted a 2.1x and 2.8x increase in infected cells (NS1+) following exposure to lower and higher STX concentrations, respectively (Figure 3D). These results mirror those obtained with cortical neurons (Figures 2A and 2D).

**Figure 3.**
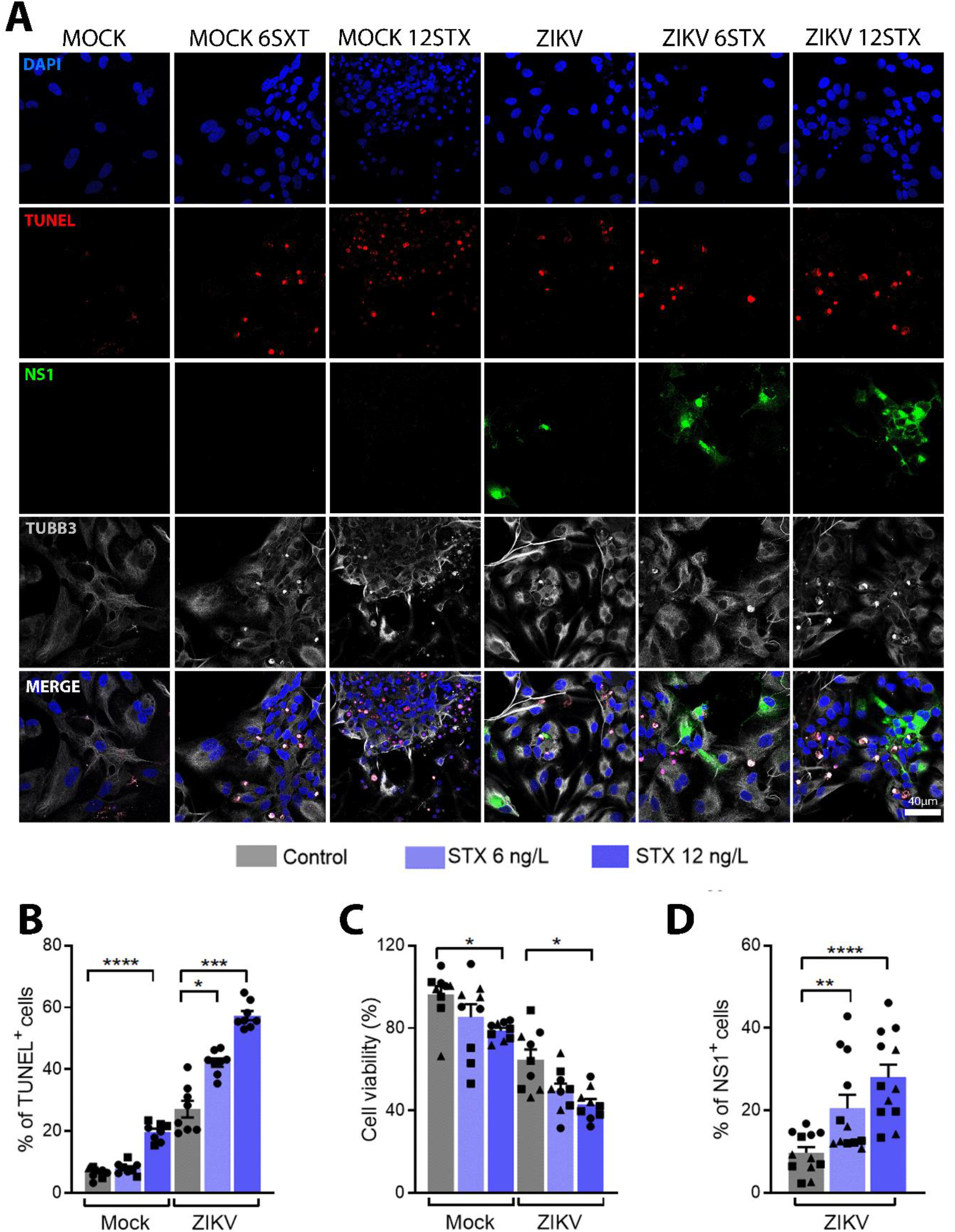
Saxitoxin amplifies the severity of ZIKV infection in hiPSC-derived sensory neurons by increasing both the rate of infection and the extent of cell death. (A) Representative photomicrographs of NS1-positive cells (green) and TUNEL-positive cells (red) of untreated or STX-treated Mock and ZIKV-infected hiPSC-derived sensory neurons. Scale bar: 40 μm. (B) Percentage of TUNEL-positive cells per DAPI-positive cells. (C) MTT cell viability assay in hiPSC-derived sensory neurons following ZIKV infection and STX treatment. (D) Percentage of NS1-positive cells per DAPI-positive cells. Experiments were performed in triplicates, in 3 independent differentiations, using 1 cell line per condition pre-treated with saxitoxin for 7 days, infected with ZIKV, and incubated with saxitoxin again for 3 days. Data is presented as average ± SEM; *p<0.05; **p<0.01; ***p<0.001 and **** p<0.0001. One-way ANOVA followed by Tukey’s multiple comparisons test was performed.

To verify the neuron-specific effects of STX, we replicated the exposure in neural progenitors and astrocytes using the highest STX concentration (12 ng/L) with ZIKV (supplementary figure 2A-2F). No changes in infection percentage (NS1+ cells) or cell viability (MTT assay) were noted in these cell types (supplementary figure 2).

## DISCUSSION

During the 2015-2016 Zika outbreak, Brazil, particularly its Northeast region, saw a disproportionately higher occurrence of congenital Zika virus syndrome (CZS) cases with severe malformations. This regional discrepancy led us to investigate potential cofactors, including socioeconomic status and environmental exposure to toxic substances. The Northeast region of Brazil, characterized by lower per capita income and poor living conditions, has previously been associated with an increased risk of ZIKV-related microcephaly, underscoring the potential influence of socioeconomic factors on disease prevalence (31,32).

Our laboratory reported that saxitoxin (STX), a toxin produced by cyanobacteria present in freshwater reservoirs in Northeast Brazil, exacerbates ZIKV-induced neurological damage in newborn mice and brain organoids. This finding suggested STX as a potential cofactor for ZIKV infection, intensifying neuropathological manifestations (14).

The present study demonstrated that STX exacerbates ZIKV-induced cell death with a marked reduction in neuronal cell types, regardless of their developmental stage. Intriguingly, while STX seemed to diminish the neuronal population, it did not further impair the effects of ZIKV on its primary target cells, namely astrocytes and neural stem cells.

Other studies have reported ZIKV’s capacity to infect and damage various types of neurons, leading to neurological disorders (33). We found that STX further exacerbates ZIKV infection and neuronal cell death, in both cortical and sensory neurons derived from human induced pluripotent stem cells (hiPSCs).

While our group previously reported that astrocytes are more affected by ZIKV than other neural cell types (16), our current study found no increase in ZIKV-induced cell death in astrocytes due to STX. Therefore, the increased severity of CZS due to concurrent ZIKV and STX exposure is likely due to a shift towards neuronal infection and subsequent damage, rather than amplified astrocyte damage.

While numerous studies underscore the specific targeting of neural progenitor cells by ZIKV, leading to brain underdevelopment (34), our findings propose a less pronounced role for neural progenitor damage when saxitoxin (STX) is introduced as a co-factor in ZIKV-induced Congenital Zika Syndrome (CZS). Contrary to expectations, we did not observe any amplification in cell death among neural progenitors when ZIKV and STX were simultaneously present.

This study explores the combined impact of water toxins and viral infections in multiple human neural cell types. The synergistic impact of toxins on viral infections is a relatively unexplored research area. Previous studies have reported increased viral replication with toxins such as STX and tetrodotoxin (TTX), emphasizing the need for further research in this field (35,36).

Our current study strengthens the evidence for STX as a cofactor amplifying the effects of ZIKV infection, predominantly in neurons. Given STX’s well-known disruptive effects on neuronal function, it is unsurprising that the combined effect of STX and ZIKV demonstrates neuronal selectivity.

Our findings underscore the importance of stringent regulations in monitoring and eliminating cyanobacterial toxins through water treatment, especially during drought periods. We found that even STX concentrations deemed safe by Brazilian authorities can enhance the effects of ZIKV. Therefore, implementing rigorous standards and surveillance of drinking water in regions with prevalent ZIKV is crucial to mitigate the harmful impact of arboviruses on human populations.

## ACKNOWLEDGMENTS

We acknowledge Pablo Trindade and Renato Molica for the helpful discussions; We recognize the contributions on technical support of Ismael Carlos da Silva Gomes, Jhonata de Sousa do Nascimento, and Beatriz Luzia De Mello Lima Guimaraes.

## AUTHOR CONTRIBUTIONS

LS, MZPG, and SKR conceived and designed the study. LS, CP, TP, CS, and GV performed all cultivations and experiments with iPSC-derived cells. LH and AT prepared and provided ZIKV. LS and MZPG interpreted data, wrote, prepared figures, and revised the manuscript. MZPG and SKR coordinated the study. All authors reviewed and contributed to the final version of this manuscript.

## ADDITIONAL INFORMATION

Competing Interests: The author(s) declare no financial or non-financial competing interests.

## SUPPLEMENTARY MATERIAL

**Supplementary Figure 1.**
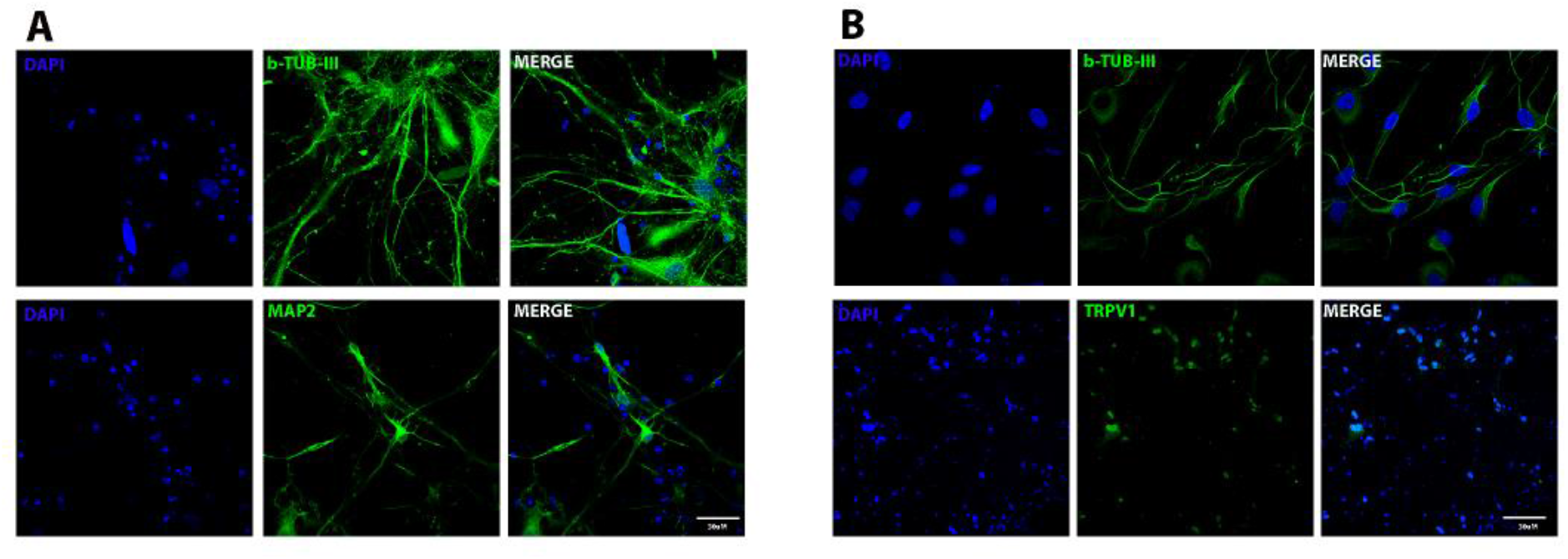
Characterization of hiPSC-derived cortical neurons. (A) Representative photomicrographs of hiPSC-derived cortical neurons stained with Beta-tubulin III and MAP2 (Green) (B) Representative photomicrographs of hiPSC-derived sensory neurons stained with Beta-tubulin III and TRPV1 (green) Scale bar: 30 μm.

**Supplementary Figure 2.**
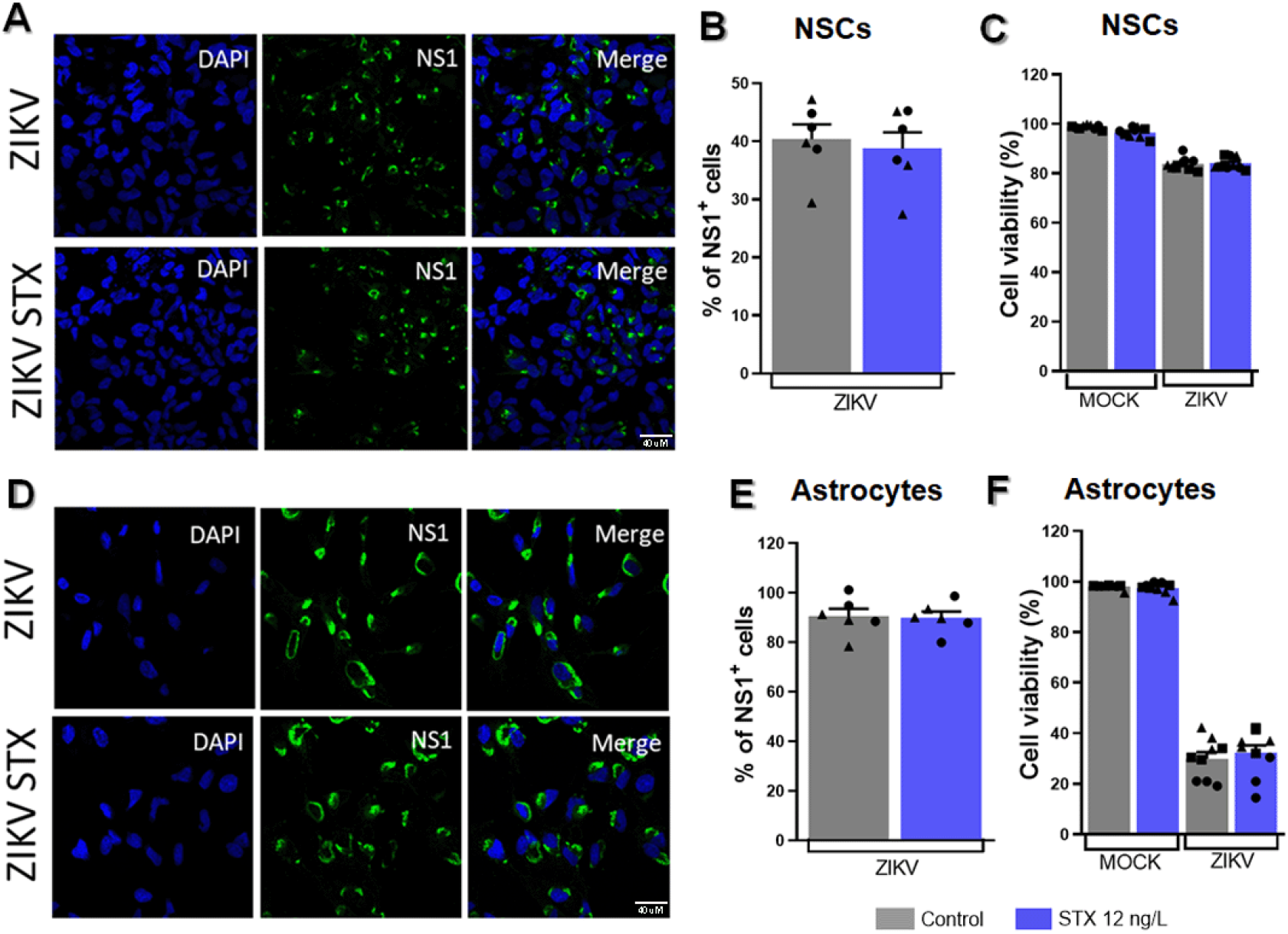
Saxitoxin does not enhance cell death of ZIKV-infected hiPSC-derived neural progenitors and astrocytes. (A) Representative photomicrographs of NS1-positive cells (green) of STX-treated Mock and ZIKV-infected hiPSC-derived neural progenitors. Scale bar: 40μm. (B) Percentage of NS1-positive cells per DAPI-positive cells.(C) MTT cell viability assay in neural progenitors following ZIKV infection and STX treatment. Experiments were performed in triplicates, in 3 independent differentiations, using 2 cell lines per condition pre-treated with saxitoxin for 7 days, infected with ZIKV, and incubated with saxitoxin again for 3 days. (D) Representative photomicrographs of NS1-positive cells (green) of STX-treated Mock and ZIKV-infected hiPSC-derived astrocytes. Scale bar: 40 μm. (E) Percentage of NS1-positive cells per DAPI-positive cells. (F) MTT cell viability assay in astrocytes following ZIKV infection and STX treatment. Experiments were performed in triplicates, in 3 independent differentiations, using 2 cell lines per condition pre-treated with saxitoxin for 7 days, infected with ZIKV, and incubated with saxitoxin again for 3 days.

## REFERENCES

1. Zanluca C, Melo VCA de, Mosimann ALP, Santos GIV dos, Santos CND dos, Luz K. First report of autochthonous transmission of Zika virus in Brazil. Mem Inst Oswaldo Cruz. 2015 Jun 9;110(4):569–72.

2. Cao-Lormeau VM, Blake A, Mons S, Lastère S, Roche C, Vanhomwegen J, et al. Guillain-Barré Syndrome outbreak associated with Zika virus infection in French Polynesia: a case-control study. The Lancet. 2016 Apr;387(10027):1531–9.

3. Rasmussen SA, Jamieson DJ, Honein MA, Petersen LR. Zika Virus and Birth Defects — Reviewing the Evidence for Causality. N Engl J Med. 2016 May 19;374(20):1981–7.

4. Nem de Oliveira Souza I, Frost PS, França JV, Nascimento-Viana JB, Neris RLS, Freitas L, et al. Acute and chronic neurological consequences of early-life Zika virus infection in mice. Sci Transl Med. 2018 Jun 6;10(444):eaar2749.

5. Aragao MFVV, Holanda AC, Brainer-Lima AM, Petribu NCL, Castillo M, van der Linden V, et al. Nonmicrocephalic Infants with Congenital Zika Syndrome Suspected Only after Neuroimaging Evaluation Compared with Those with Microcephaly at Birth and Postnatally: How Large Is the Zika Virus “Iceberg”? AJNR Am J Neuroradiol. 2017 Jul;38(7):1427–34.

6. Driggers RW, Ho CY, Korhonen EM, Kuivanen S, Jääskeläinen AJ, Smura T, et al. Zika Virus Infection With Prolonged Maternal Viremia and Fetal Brain Abnormalities. Obstetric Anesthesia Digest. 2017 Mar;37(1):51–51.

7. Campos GS, Bandeira AC, Sardi SI. Zika Virus Outbreak, Bahia, Brazil. Emerg Infect Dis. 2015 Oct;21(10):1885–6.

8. Molica RJR, Oliveira EJA, Carvalho PVVC, Costa ANSF, Cunha MCC, Melo GL, et al. Occurrence of saxitoxins and an anatoxin-a(s)-like anticholinesterase in a Brazilian drinking water supply. Harmful Algae. 2005 Jun;4(4):743–53.

9. Subramanian N, Wetzel A, Dombert B, Yadav P, Havlicek S, Jablonka S, et al. Role of Nav1.9 in activity-dependent axon growth in motoneurons. Human Molecular Genetics. 2012 Aug 15;21(16):3655–67.

10. Kao CY, Nishiyama A. Actions of saxitoxin on peripheral neuromuscular systems. J Physiol. 1965 Sep;180(1):50–66.

11. Zakon HH. Adaptive evolution of voltage-gated sodium channels: The first 800 million years. Proc Natl Acad Sci USA. 2012 Jun 26;109(supplement_1):10619–25.

12. Rufino R, Gracie R, Sena A, Freitas CM de, Barcellos C. Surtos de diarreia na região Nordeste do Brasil em 2013, segundo a mídia e sistemas de informação de saúde – Vigilância de situações climáticas de risco e emergências em saúde. Ciênc saúde coletiva. 2016 Mar;21(3):777–88.

13. De Araújo TVB, Ximenes RADA, Miranda-Filho DDB, Souza WV, Montarroyos UR, De Melo APL, et al. Association between microcephaly, Zika virus infection, and other risk factors in Brazil: final report of a case-control study. The Lancet Infectious Diseases. 2018 Mar;18(3):328–36.

14. Pedrosa C da SG, Souza LRQ, Gomes TA, de Lima CVF, Ledur PF, Karmirian K, et al. The cyanobacterial saxitoxin exacerbates neural cell death and brain malformations induced by Zika virus. Pimenta PFP, editor. PLoS Negl Trop Dis. 2020 Mar 12;14(3):e0008060.

15. Sochacki J, Devalle S, Reis M, Mattos P, Rehen S. Generation of urine iPS cell lines from patients with Attention Deficit Hyperactivity Disorder (ADHD) using a nonintegrative method. Stem Cell Research. 2016 Jul;17(1):102–6.

16. Ledur PF, Karmirian K, Pedrosa C da SG, Souza LRQ, Assis-de-Lemos G, Martins TM, et al. Zika virus infection leads to mitochondrial failure, oxidative stress and DNA damage in human iPSC-derived astrocytes. Sci Rep. 2020 Jan 27;10(1):1218.

17. Garcez PP, Loiola EC, Madeiro da Costa R, Higa LM, Trindade P, Delvecchio R, et al. Zika virus impairs growth in human neurospheres and brain organoids. Science. 2016 May 13;352(6287):816–8.

18. Pedrosa C da SG, Goto-Silva L, Temerozo JR, Souza LRQ, Vitória G, Ornelas IM, et al. Non-permissive SARS-CoV-2 infection in human neurospheres. Stem Cell Research. 2021 Jul;54:102436.

19. Casas BS, Vitória G, Prieto CP, Casas M, Chacón C, Uhrig M, et al. Schizophrenia-derived hiPSC brain microvascular endothelial-like cells show impairments in angiogenesis and blood–brain barrier function. Mol Psychiatry. 2022 Sep;27(9):3708–18.

20. Casas BS, Vitória G, do Costa MN, Madeiro da Costa R, Trindade P, Maciel R, et al. hiPSC-derived neural stem cells from patients with schizophrenia induce an impaired angiogenesis. Transl Psychiatry. 2018 Feb 22;8(1):48.

21. Takahashi K, Tanabe K, Ohnuki M, Narita M, Ichisaka T, Tomoda K, et al. Induction of Pluripotent Stem Cells from Adult Human Fibroblasts by Defined Factors. Cell. 2007 Nov;131(5):861–72.

22. Yan Y, Shin S, Jha BS, Liu Q, Sheng J, Li F, et al. Efficient and Rapid Derivation of Primitive Neural Stem Cells and Generation of Brain Subtype Neurons From Human Pluripotent Stem Cells. Stem Cells Translational Medicine. 2013 Nov 1;2(11):862–70.

23. Trindade P, Loiola EC, Gasparotto J, Ribeiro CT, Cardozo PL, Devalle S, et al. Short and long TNF-alpha exposure recapitulates canonical astrogliosis events in human-induced pluripotent stem cells-derived astrocytes. Glia. 2020 Jul;68(7):1396–409.

24. Guimarães MZP, De Vecchi R, Vitória G, Sochacki JK, Paulsen BS, Lima I, et al. Generation of iPSC-Derived Human Peripheral Sensory Neurons Releasing Substance P Elicited by TRPV1 Agonists. Front Mol Neurosci. 2018 Aug 22;11:277.

25. Goto-Silva L, Ayad NME, Herzog IL, Silva NP, Lamien B, Orlande HRB, et al. Computational fluid dynamic analysis of physical forces playing a role in brain organoid cultures in two different multiplex platforms. BMC Dev Biol. 2019 Dec;19(1):3.

26. Lancaster MA, Knoblich JA. Generation of cerebral organoids from human pluripotent stem cells. Nat Protoc. 2014 Oct;9(10):2329–40.

27. Herculano-Houzel S, Lent R. Isotropic Fractionator: A Simple, Rapid Method for the Quantification of Total Cell and Neuron Numbers in the Brain. J Neurosci. 2005 Mar 9;25(10):2518–21.

28. Carod-Artal FJ. Neurological complications of Zika virus infection. Expert Review of Anti-infective Therapy. 2018 May 4;16(5):399–410.

29. Martinez ARM, Costa MCM, Novaes MAC, Lima HC, Nucci A, França MC. A novel phenotype Of Zika virus-related neurological disease: Sensory neuronopathy: Noteworthy Cases. Muscle Nerve. 2018 Feb;57(2):E100–1.

30. Oh Y, Zhang F, Wang Y, Lee EM, Choi IY, Lim H, et al. Zika virus directly infects peripheral neurons and induces cell death. Nat Neurosci. 2017 Sep 1;20(9):1209–12.

31. Barbeito-Andrés J, Schuler-Faccini L, Garcez PP. Why is congenital Zika syndrome asymmetrically distributed among human populations? Riley S, editor. PLoS Biol. 2018 Aug 24;16(8):e2006592.

32. Souza WVD, Albuquerque MDFPMD, Vazquez E, Bezerra LCA, Mendes ADCG, Lyra TM, et al. Microcephaly epidemic related to the Zika virus and living conditions in Recife, Northeast Brazil. BMC Public Health. 2018 Dec;18(1):130.

33. Swartwout B, Zlotnick M, Saver A, McKenna C, Bertke A. Zika Virus Persistently and Productively Infects Primary Adult Sensory Neurons In Vitro. Pathogens. 2017 Oct 13;6(4):49.

34. Wen Z, Song H, Ming G li. How does Zika virus cause microcephaly? Genes Dev. 2017 May 1;31(9):849–61.

35. Bogomolni AL, Bass AL, Fire S, Jasperse L, Levin M, Nielsen O, et al. Saxitoxin increases phocine distemper virus replication upon in-vitro infection in harbor seal immune cells. Harmful Algae. 2016 Jan;51:89–96.

36. Zhang CX, Ofiyai H, He M, Bu X, Wen Y, Jia W. Neuronal activity regulates viral replication of herpes simplex virus type 1 in the nervous system. J Neurovirol. 2005 Jan;11(3):256–64.

